# Human cartilage progenitor cells from ear, nose, rib, and joint have a robust, stable phenotype for cartilage repair

**DOI:** 10.1101/2022.06.30.498323

**Authors:** Soheila Ali Akbari Ghavimi, Paul M Gehret, Terri Giordano, Kyra W Y Smith, Riccardo Gottardi

## Abstract

**Background:** Cartilage progenitor cells (CPCs) are a small but highly proliferative cell population that resides within cartilage. Joint cartilage CPCs have a high chondrogenic potential and superior cartilage formation characteristics; however, CPCs from other cartilage sources more accessible for translation such as ear, nose, and rib are broadly unexplored. Our study illuminates the differences between CPCs from these four cartilages, their corresponding tissue chondrocyte (CC), and bone marrow-derived mesenchymal stem cell (MSC).

**Methods:** CPCs subtypes were isolated from pediatric cartilage via fibronectin selection, immunophenotyped by flow cytometry and compared to MSCs. Trilineage differentiation capacity was assessed via histology and qRT-PCR. Next, triiodothyronine was used to hypertrophically challenge each CPC subset and their corresponding chondrocyte population. After 28 days cartilage pellets were assessed via histology, immunohistochemistry, and qRT-PCR.

**Findings:** Each CPC subset possessed a specific immunophenotypic signature with CD56 as a potential common marker. All CPC subsets proliferated 2-fold faster than MSCs and 4-fold faster than CCs. Additionally, CPCs had a substantially reduced propensity for osteogenic differentiation and very limited adipogenic capacity by histology and gene expression. Finally, all CPC subsets resisted the hypertrophic challenge more than the corresponding chondrocyte population marked by less collagen X secretion and downregulation of hypertrophy associated genes.

**Interpretation:** CPCs represent a promising cell type for cartilage regeneration. The ease of accessibility of the ear and nose CPCs present opportunities for new translational approaches and reduced clinical timelines.

**Funding:** CHOP Research Institute, Frontier Program in Airway Disorders of CHOP, NIH (R21HL159521), NSF-GRFP (DGE-1845298)

## 1. Introduction

Cartilage progenitor cells (CPCs), recognized by their colony forming capacity and rapid self-renewal, have been shown to subside in small subpopulations within articular cartilage.^1,2^ At first cartilage progenitors were associated with the synovium as a source of repair for joint diseased states like osteoarthritis;^1^ however, CPC populations have progressively been recognized to be present also within healthy cartilage using *in vivo* cell proliferation toolkits such as bromodeoxyuridine labeling.^3,4^ CPCs have unique attributes such as a minimal osteogenic and adipogenic capacity, longer telomere lengths, and different surface markers when compared to mesenchymal stem cells (MSCs) and chondrocytes (CCs).^2,5,6^ CPC’s chondrogenic priming, low propensity to calcify, and rapid proliferation rate have made them prime candidates for cartilage repair in the joint.^7^ However, while articular CPCs are important for understanding joint development and regeneration, cellular therapies using joint CPCs are faced by the major hurdle of extracting the cells from the joint, in itself a very invasive procedure that poses significant challenges to translation.

Given the relative inaccessibility of joint CPCs, the field of cartilage repair has been more focused on MSCs as a potentially more feasible approach. MSCs have good proliferation, trilineage differentiation capacity, and are routinely harvested in the clinic from iliac crest biopsies making them a more readily available cell type compared to joint CPCs. For over two decades MSCs had been hailed as the future of cartilage repair.^8^ However, MSCs’ suffer from significant donor variability^9–11^ and have a tendency to undergo hypertrophy^12^ with a high associated risk of calcification.^13,14^ This has resulted in the progressive decline of clinical trials utilizing MSCs, and out of the 914 phase 2 trials conducted over the past 2 decades, only 6% have progressed successfully to phase 3.^15,16^ This somewhat limited translational success has a resulted in a decreased enthusiasm for MSCs. Consequently, over the past few years the interest in leveraging CPCs for cartilage repair has been rapidly increasing, focusing on the possibility of accessing alternative cartilage sources besides the joint such as auricular,^17–22^ nasal,^7^ and costal^23^ cartilage.

Biopsies and grafts from each of these cartilage types are routinely harvested in the clinic and represent a less invasive and more practical approach to obtain CPCs for cartilage repair.^24–27^ Among these options, ear CPCs (eCPCs) are the easiest to harvested through a minimally invasive biopsy, a procedure that is commonly perform for tympanoplasty, is painless and leaves no visible scar.^24^ Human eCPCs have been recently isolated for auricular tissue engineering by Otto et al.^17^ Furthermore, eCPCs have been isolated and characterized to a limited extent from lapine,^21^ porcine,^18,19^ cercopithecine,^20^ and canine ^22^ models. Similarly to ears, nasal cartilage offers relative ease of access for biopsy and no donor site morbidity compared to other joint cartilages,^28^ but have been less explored.

Jessop et. al. is the only group who has isolated and characterized nose CPCs (nCPCs) derived from human nasoseptal cartilage.^7^ Finally, rib cartilage is one of the most common sources of autologous grafts due to its abundance and mechanical strength allowing surgeons to harvest from a single donor with fewer complications.^29^ However, rib CPCs have barely been studied.

Cartilage tissues from different sites have different characteristics as shown in Figure 1, where histological and immunohistochemical staining of pediatric ear, nose, rib, and joint (digit) cartilage are compared to highlight the differences between matrix composition, structure, cell density and cell organization. Each type of cartilage presents a specific morphology, and this clearly raises the question of how similar the CPC populations within each of these cartilage types are to not only joint CPCs but also to MSCs and to autologous CCs. The goal of the present study is to evaluate if human CPCs harvested from these different anatomic locations present different surface markers, proliferation ability, differentiation capacity, matrix secretion profile, and hypertrophic resistance compared to both MSCs and CCs.

**Figure 1.**
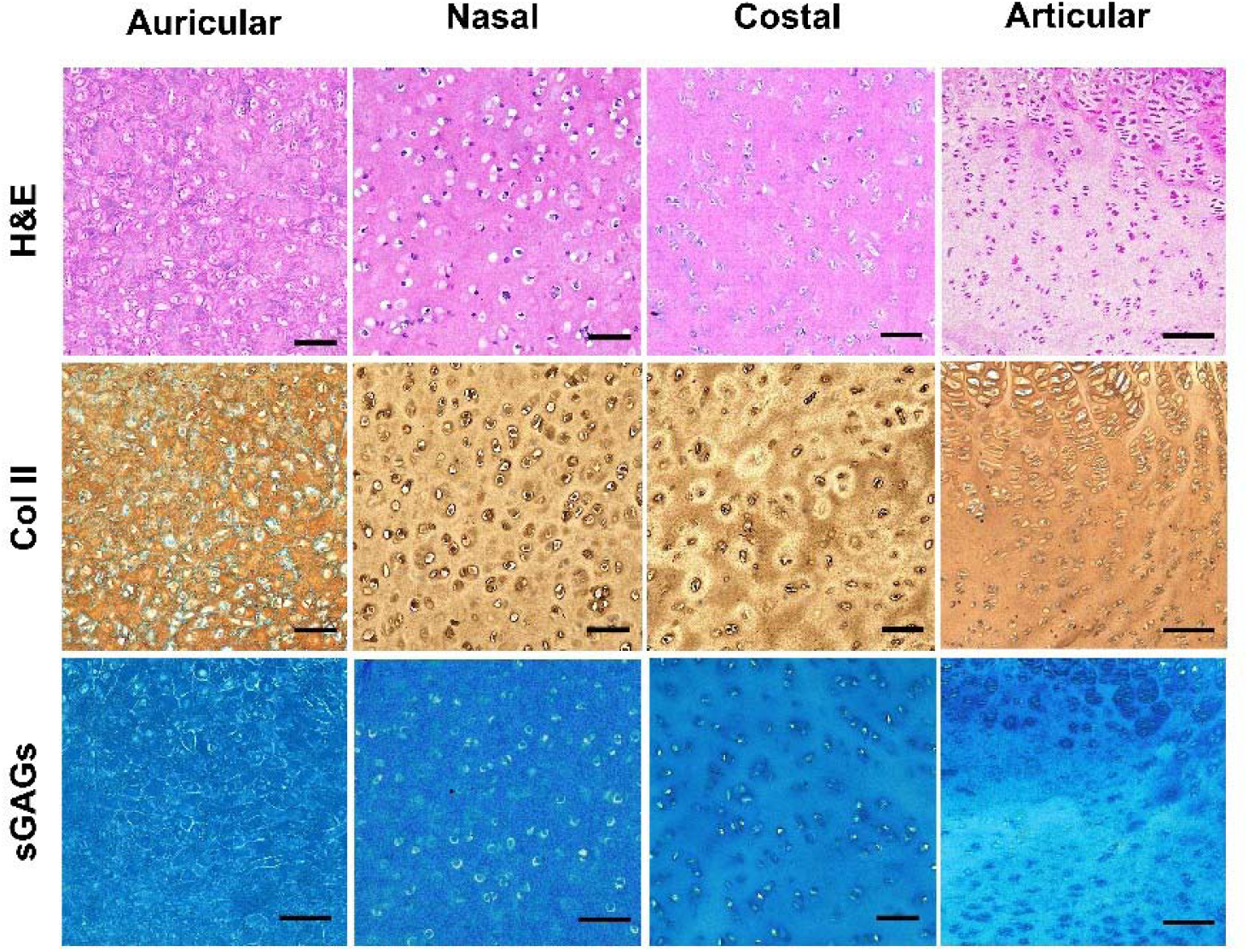
Composition and structure of the pediatric cartilage utilized in this study. Auricular, nasal, costal, and articular cartilage stained using H&E (row 1), IHC for Col II (row 2), or Alcian blue (row 3).

## 2. Materials and Methods

### 2.1. CPC Isolation and Expansion

Human cartilage tissue from the ear, nose, rib and supernumerary digits (n=4 for each tissue type, total of n=16) was collected from the surgical waste of pediatric patients (aged 0 to 18 years old) undergoing tympanoplasty, rhinoplasty, airway reconstruction, or supernumerary digits excision, respectively (Children’s Hospital of Philadelphia IRB 20-017817, 20-017551, 20-017287). Cells were extracted by sequential pronase (70 U/mL, 30 min, 37ºC; Sigma) and collagenase II (300 U/ml, 4 h, 37ºC; Worthington Biochemical) digestion. Then, cartilage-resident progenitor cells (CPCs) were separated from cartilage chondrocytes (CCs) by selective fibronectin-adhesion as originally described in Dowthwaite *et al*.^18^ and Williams *et al*.^30^ Briefly, 1000 cells/well were seeded on fibronectin-coated 6-well plates at 37°C. After 20 mins, non-adherent cells (CCs) were removed and expanded in new plates in Dulbecco’s Modified Eagle Medium (DMEM; Gibco) containing 10% Fetal Bovine Serum (FBS; Gibco) and 2% Penicillin-Streptomycin-Fungizone (PSF, Anti-Anti; Gibco) until passage 4. Adherent cells (CPCs) were cultured in DMEM/F12 (Gibco) with 10% FBS, 2% FBS, 10 mM HEPES, 0.5 mg/ml L-Glucose, and 0.1 mM L-ascorbic acid 2-phosphate. CPCs formed colonies and reached confluency after ∼12 days, after which they were expanded until passage 4 in the same medium with the addition of 5 ng/ml FGF-2, and 1 ng/ml TGF-β1. Bone-marrow derived mesenchymal stem cells (MSCs) (n=4 donors) were purchased from Roosterbio at passage 2 and expanded in Roosterbio growth media until passage 4.

### 2.2. Growth Curve

To compare the proliferation rate of CPCs, CCs, and MSCs cells at passage 4 were seeded in 150 cm^2^ culture flasks at the concentration of 2×10^5^ cells/flask and grown in their respective growth medium. Cells were harvested at day 1, 2, 3, 5, 7, 9 then counted to plot as a growth curve.

### 2.3. Flow cytometry

CPCs and MSCs at passage 3 were suspended in cell staining buffer (Biolegend) (10^6^ cells/100μl), pre-incubated and blocked with 5μl of Human Fc Receptor Blocking Solution (Biolegend) for 10 mins at room temperature then stained with 5μl/10^6^ cells fluorescent conjugated Brilliant Violet 510™ anti-human CD14, Brilliant Violet 605™ anti-human CD19, PerCP/Cy5.5 anti-human CD29, Brilliant Violet 650™ anti-human CD34, FITC anti-human CD44, PE/Cy7 anti-human CD56 (NCAM), PE anti-human CD73, APC anti-human CD45, APC/Cyanine7 anti-human CD90, and Brilliant Violet 421™ anti-human CD105 (Biolegend) for 20 min on ice. Cells were washed and resuspended in 500μl of staining buffer. Unstained cells were used as negative control and UltraComp eBeads® (Invitrogen) were used as single-color compensation controls to determine appropriate FSC/SSC settings. A minimum of 10,000 events were collected for each sample using Cytek Aurora and analyzed by FCS Express™. The positivity percentage of each surface marker was determined using the subtraction plot where the percentage of positive cells was quantified.

### 2.4. Tri-lineage differentiations

CPCs and MSCs were seeded in 6-well plate at 10^6^ cells/well and treated with either osteogenic (DMEM, 10% FBS, 2% FBS, 0.1 μM dexamethasone, 10 mM β-glycerophosphate, 50 μg/ml L-ascorbic acid 2-phosphate, 10 nM 1α, 25-Dihydroxyvitamin D3), chondrogenic (DMEM, 2% FBS, 10 μg/ml Insulin-Transferrin-Selenium, 0.1 μM Dexamethasone, 40 μg/mL L-proline, 50 μg/mL L-ascorbic acid 2-phosphate, and 10 ng/mL TGF-β3 (PeproTech)), or adipogenic (DMEM, 10% FBS, 2% FBS, 1 μg/ml Insulin-Transferrin-Selenium, 0.1 μM Dexamethasone, and 10 mM 3-isobutyl-1-methylxantine (IBMX; Sigma)) media renewed every 3 days. To form pellets, CPCs and MSCs were centrifuged in a v-bottom 96-well plate to form pellets that were cultured in chondrogenic medium renewed every 3 days. After 21 days, cells in wells and pellets were washed and fixed with 10% formalin. Cells in plates were stained Alizarin Red, Alcian Blue, or Oil Red O corresponding to their stimulation with either osteogenic, chondrogenic, or adipogenic media, respectively, as previously described.^31–35^ Pellets were paraffin embedded and sectioned (see immunohistochemistry section) prior to Alcian Blue staining. All samples were imaged with BZ-X810 All-in-One Microscope (Keyence). Replicates for all samples were treated with Trizol to extract RNA for gene expression analysis (see qRT-PCR section).

### 2.5. Chondrogenic differentiation and enhancement of hypertrophy

CPCs, CCs, and MSCs were seeded (2×10^5^ cells/well) in a V-bottom 96 well plate, centrifuged at 250 g for 5 min, and exposed to chondrogenic medium for 14 days followed by another 14 days of either chondrogenic or hypertrophic medium (DMEM, 2% FBS, 10 μg/ml Insulin-Transferrin-Selenium, 40 μg/mL L-proline, 50 μg/mL L-ascorbic acid 2-phosphate, and 1 nM triiodothyronine (T3; Sigma)). After 28 days, pellets were washed and collected for immunohistochemistry staining, biochemical assays, and RT-PCR.

### 2.6. Immunohistochemical staining

Pellets of CPCs, CCs, and MSCs were collected at day 14 and 28, washed with PBS, then fixed in 10% formalin overnight at 4 °C. Next the pellets were dehydrated (70% isopropanol, 96% isopropanol, 100% isopropanol, 100% acetone, pre-warmed paraffin) and embedded in paraffin. The paraffin blocks were sectioned, deparaffinized (heat and histoclear) and rehydrated by inverse ethanol dilutions and water. After antigen retrieval with chondroitinase ABC (100 mU/ml) and hyaluronidase (250 U/ml) in 0.02 % BSA for 30 min at 37 °C, the tissue sections were stained for collagen I (MA126771; Life Technologies Corporation), collagen II (MA512789; Thermofisher), or collagen X (14-9771-82; Invitrogen) then visualized using a mouse and rabbit specific HRP/DAB (ABC) detection IHC kit (ab64264) as per the manufacturer’s protocol. Stained pellets were counterstained with hematoxylin prior to visualization by the Zeiss Slide Scanner.

### 2.7. Biochemical assays

This protocol was adapted from Markway et. al.^36^ Pellet of CPCs, CCs, and MSCs were collected at day 14 and 28, digested for 24 hours at 60ºC in 125 μg/mL papain in 5 mM EDTA and 5 mM L-cysteine at 60°C followed by further mechanical disrupted by pellet pestles. Following digestion either 1,9 dimethylmethylene blue (DMMB) or PicoGreen™ was added to the samples to quantify sulfated glycosaminoglycans or DNA respectively. The total GAG or DNA content was calculated through comparison with a standard curve of chondroitin-6-sulfate or lambda phage DNA. The same digestates were mixed with 10 N HCL 1:1 and the supernatant collected and evaporated. The samples resuspended in 1X assay buffer and mixed with 5% (w/v) chloramine T for 20 min at room temperature. Following oxidation, orthohydroxyproline (OHP, ratio 1:7) was added to the samples to quantify collagen content. The total collagen content was calculated through comparison with a standard curve of Trans-4-hydroxy-L-proline.

### 2.8. qRT-PCR

Cells differentiated in 6 well plates were treated with Trizol and the solutions were stored at -80°C until RNA extraction. Pellet samples at day 14 and day 28 were washed with PBS, and saved in RNA-Later at -80°C. For total RNA extraction, pellets were washed in PBS, flashed frozen, pulverized, and resuspended in Trizol. All Trizol solutions were then mixed with chloroform, and the RNA, DNA, and the organic layer were separated by centrifugation. RNA was purified using RNeasy Plus mini kit (Qiagen) following the manufacturer’s protocol. qRT-PCR was performed using the BioMark system (Fluidigm) according to the two-step single-cell gene expression protocol with EvaGreen as described in the Real-Time PCR analysis User Guide (PN 6800088, Fluidigm). For this procedure, 666 ng of starting RNA for each sample underwent 18 cycles of specific target amplification (STA) followed by a ten-fold dilution. STA CT values were normalized to the housekeeping genes ACTB, GAPDH, and eight genes that show the most stable expression in the SCN according to sequencing data. expression level was calculated by the 2^-ΔΔCT^ method. The expression level at day 0 for the Tri-Lineage differentiation test, at day 14 for the hypertrophic/chondrogenic pellet test, and 18S rRNA and β-actin as housekeeping genes were used as reference.

### 2.9. Statistical Analysis

All data resulted from the average of n=4 different biological replicates from 4 different human donors. Furthermore, each biological replicate measurement resulted from the average of n=5 technical replicates for IHC, biochemical assays, and qRT-PCR; and n=2 for flow cytometry, Tri-Lineage differentiation, and growth curve. A two-way ANOVA with Tukey’s *post hoc* analysis (multiple comparison) was used to calculate the statistical differences among the biological replicates. A paired, two-tailed t-test was used to evaluate statistical differences between the chondrogenic and hypertrophic treatments of same cell type. Statistical significance was established for *p* values <0.05.

## 3. Results

### 3.1. CPCs display different surface marker expression

The immunophenotypic differences between each CPC subtype compared to MSCs was obtained by flow cytometry of adherent cells at passage 3 after lifting. From the overton subtraction plot of each surface marker the percentage positive was automatically calculated for each individual donor (Figure 2). The individual histograms are reported in the supplemental (Figure S1). MSCs are known to be CD29^+^, CD44^+^, CD73^+^, CD90^+^, CD105^+^, and CD14^-^, CD19^-^, CD34^-^, CD45^-^, CD56^-^,^37^ and our results confirmed the same trend. Each CPC type presented substantial differences from the common MSC surface markers. CPCs from each tissue source were positive for CD29, CD44, CD73, and CD90, and negative for CD14, similarly to our observed MSCs immunophenotype. All other surface markers differed in some respect from MSCs depending on CPC type. CD105 is the surface receptor for transforming growth factor (TGF)-beta and MSCs, eCPCs, and nCPCs were all positive for this marker while rCPCs and dCPCs were negative.^38^ The expression of CD105 has been shown to correlate with spontaneously occurring chondrogenic differentiation and to mark populations with better chondrogenesis.^39,40^ CD19 is a receptor present in specific subsets of hematopoietic stem cells;^41^ MSCs and nCPCs were negative for this marker while eCPCs, rCPCs, and dCPCs were positive. Notably CD19 has recently been identified as surface marker of brain mural cell lineages ^42^ including pericytes, a multipotent cell type capable of robust chondrogenesis.^43^ CD34 is another, less common, hematopoietic stem cell marker and only nCPCs displayed positivity.^44^ CD45 is a surface marker commonly expressed in immune cells and most cells, as expected, were negative for this marker with the exception of eCPCs. However, recently, CD45 has also been reported as a surface marker for progenitor cells and has even been found as a marker from CPCs derived from iPSCs.^45^ Finally, CD56 is a clonal stem cell marker and all CPCs displayed positivity for this surface marker while MSCs did not.

**Figure 2.**
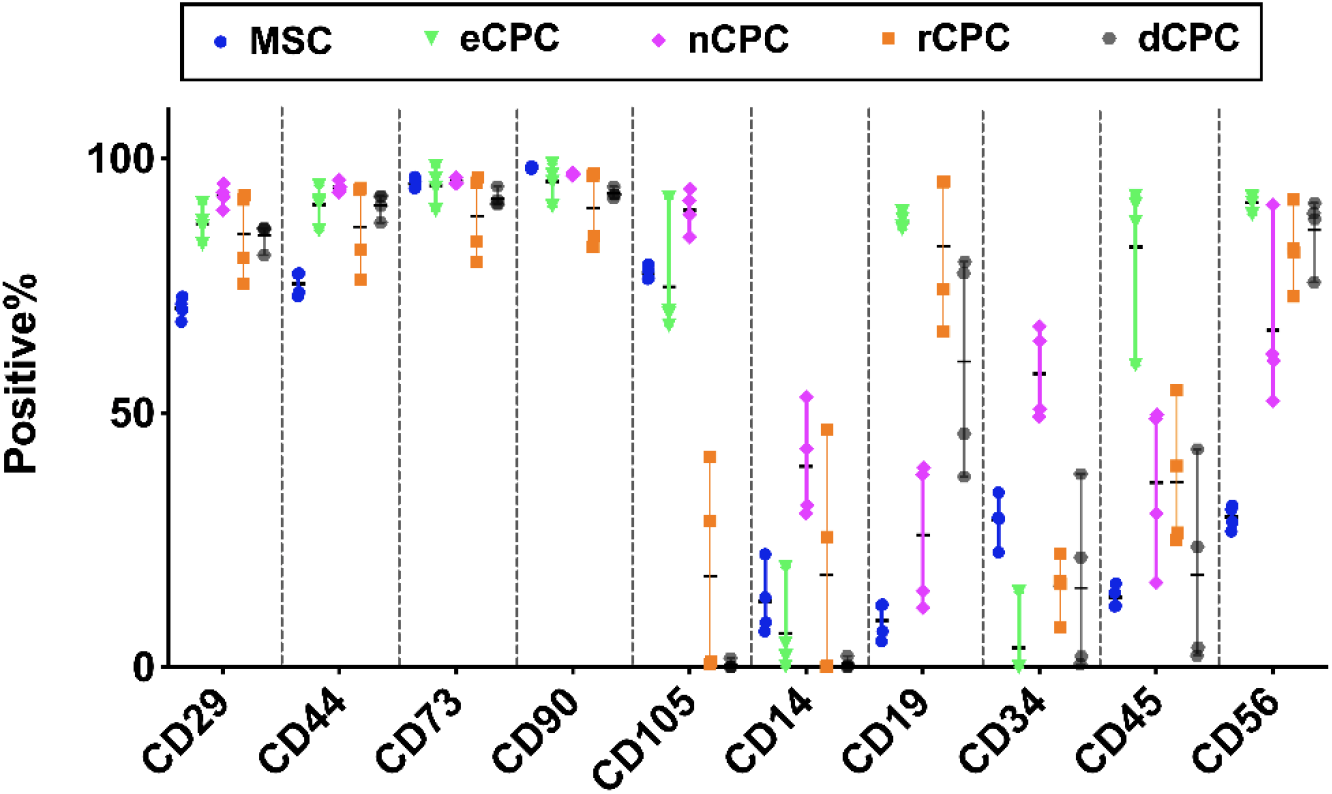
Immunophenotypic characterization of MSCs and CPCs surface markers by flow cytometry. Percent positivity of CD29, CD44, CD73, CD90, CD105, CD14, CD19, CD34, CD45, CD56 at passage 3 compared to the corresponding unstained samples. Results reported as mean ± standard deviation.

### 3.2. CPCs proliferate faster than MSCs and CCs

CPCs, MSCs, and CCs proliferation rates were assessed for 10 days *in vitro*. All CPCs subsets grew at a higher rate than both MSCs and CCs, and this more rapid growth was already evident by day 3, fully reaching confluency by day 7 (Figure 3). Growth rates were exponential for all cells until reaching confluency which occurred only for CPCs (day 7) in the timeframe considered. Overall, the digit CPCs presented the fastest proliferation rate among all CPCs followed by rib, ear, and nose. Interestingly, this same pattern was also mirrored by the corresponding chondrocyte populations.

**Figure 3.**
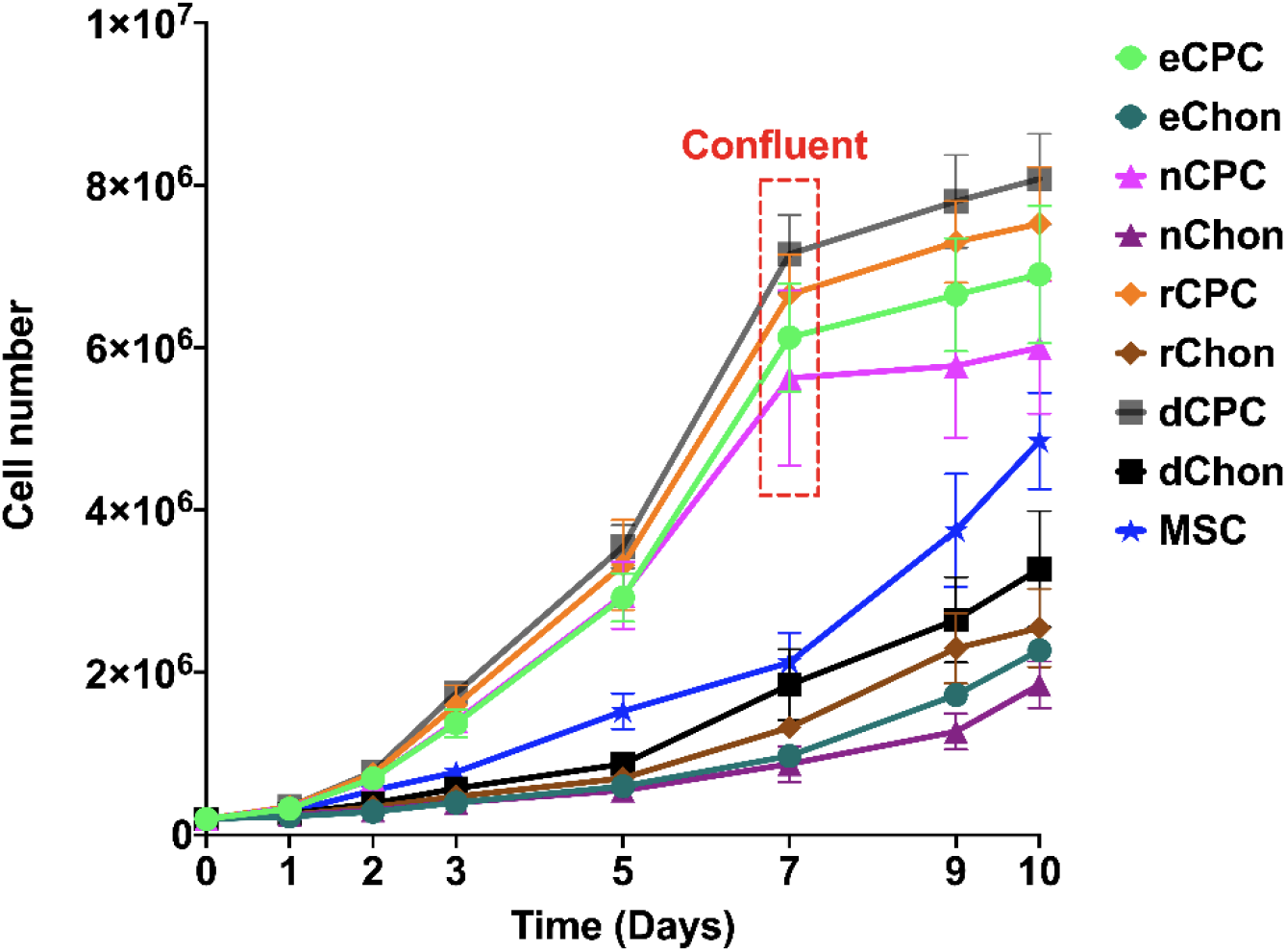
Growth curve of CPCs, MSCs and CCs. The proliferation rate of MSCs, CPCs, and the corresponding CCs over 10 days.

### 3.3. CPCs have limited osteogenic and adipogenic differentiation capacity

CPCs and MSCs differentiation potential was tested by exposing each cell type to either osteogenic, adipogenic, or chondrogenic medium. Cell pellets are the standard for evaluating chondrogenesis, thus this method was utilized to test chondrogenic potential^46^ while 2D differentiation was used for osteogenesis and adipogeneis. Representative images of MSCs and CPCs (stained with Alizarin red, Alcian blue, or Oil Red-O) indicate the cells’ capacity to form deposits of calcium phosphate, secrete glycosaminoglycans (GAGs), and develop fat droplets, respectively. Upon stimulation, MSCs showed a high aptitude to differentiate into each lineage (figure 4). As expected, all CPCs subsets showed strong differentiation toward the chondrogenic lineage. The more intense GAGs staining of the eCPCs and nCPCs pellets compared to the MSCs’ suggests that both possess a superior chondrogenic potential, likely also higher than dCPCs and rCPCs. Moreover, Alizarin red staining indicates that all CPCs showed significantly lesser propensity for osteogenesis than MSCs, although with varying degrees within the CPCs subsets. Finally, CPCs were unable to produce any fat droplets indicating minimal to no adipogenic potential.

**Figure 4.**
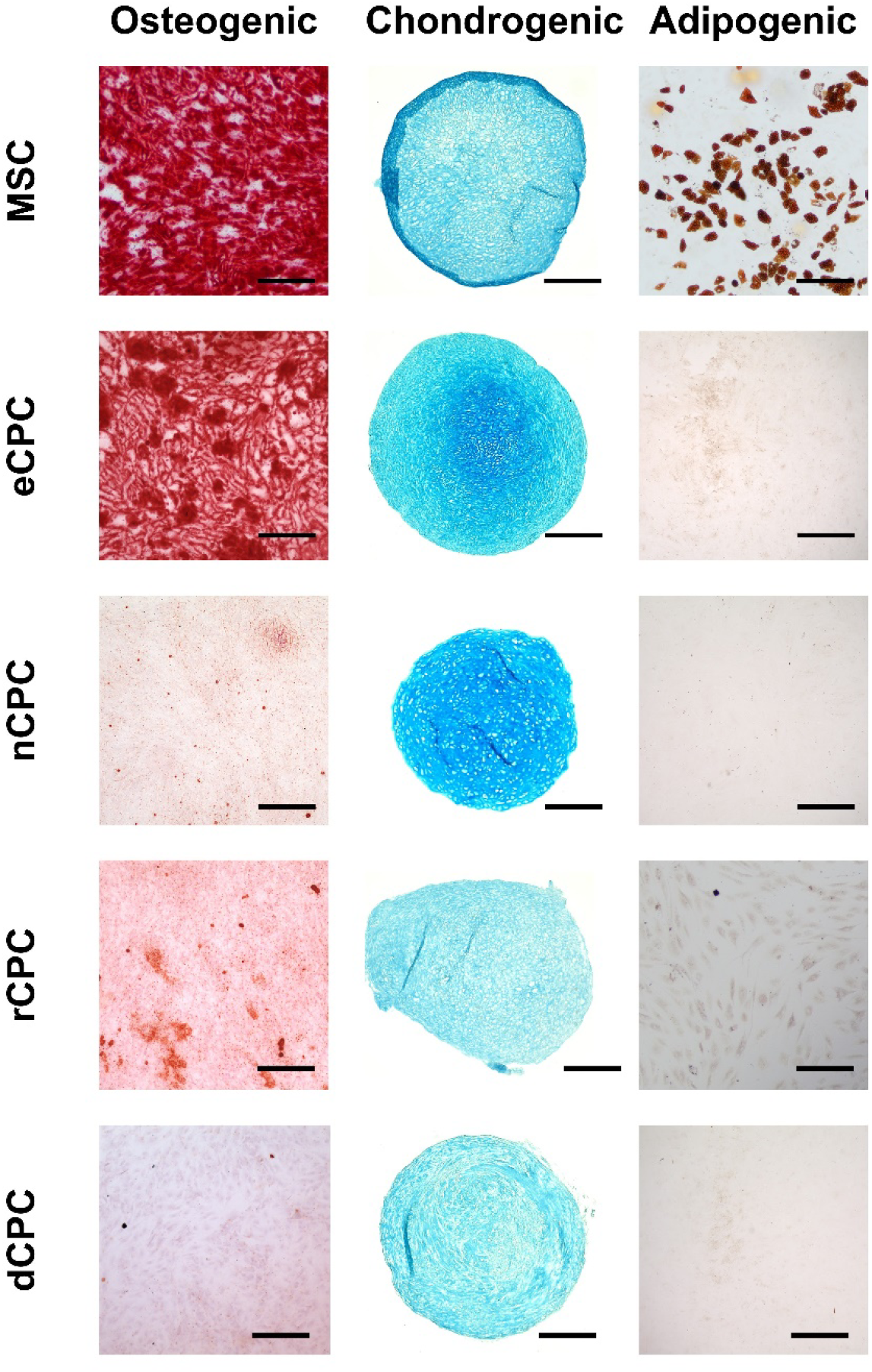
Tri-lineage differentiation of the MSCs and CPCs using osteogenic, chondrogenic, or adipogenic medium for 3 weeks. Alizarin red staining of each cell type cultured in osteogenic medium (column 1). Alcian blue staining of each cell type pellet cultured in chondrogenic medium (column 2). Oil Red-O staining of each cell type cultured in adipogenic medium (column 3). Scale bars = 100 μm.

To further explore differences in tri-lineage differentiation potential between MSCs and CPCs, gene expression was quantified for osteogenic (ALP, OPN, OCN, RUNX2, and COL1a1), chondrogenic (ACAN, SOX9, PRG4, COL10a1, COL2a1), and adipogenic (ADIPQ, PPAR□, LPL, LEP) genes (Figure 5). Osteogenically differentiated MSCs showed markedly higher expression of all osteogenic genes compared to eCPCs, nCPCs, rCPCs, and dCPCs. Among the CPC subsets, varying levels of osteogenic gene expression were observed among all CPC subsets, with eCPCs presenting the higher expression for ALP, OPN, OCN, and RUNX2, and nCPCs the lower expression levels for ALP and to a lesser extent for OCN and RUNX2. The quantitative osteogenic gene expression analysis matches the trend observed by Alizarin red staining in Figure 4.

**Figure 5.**
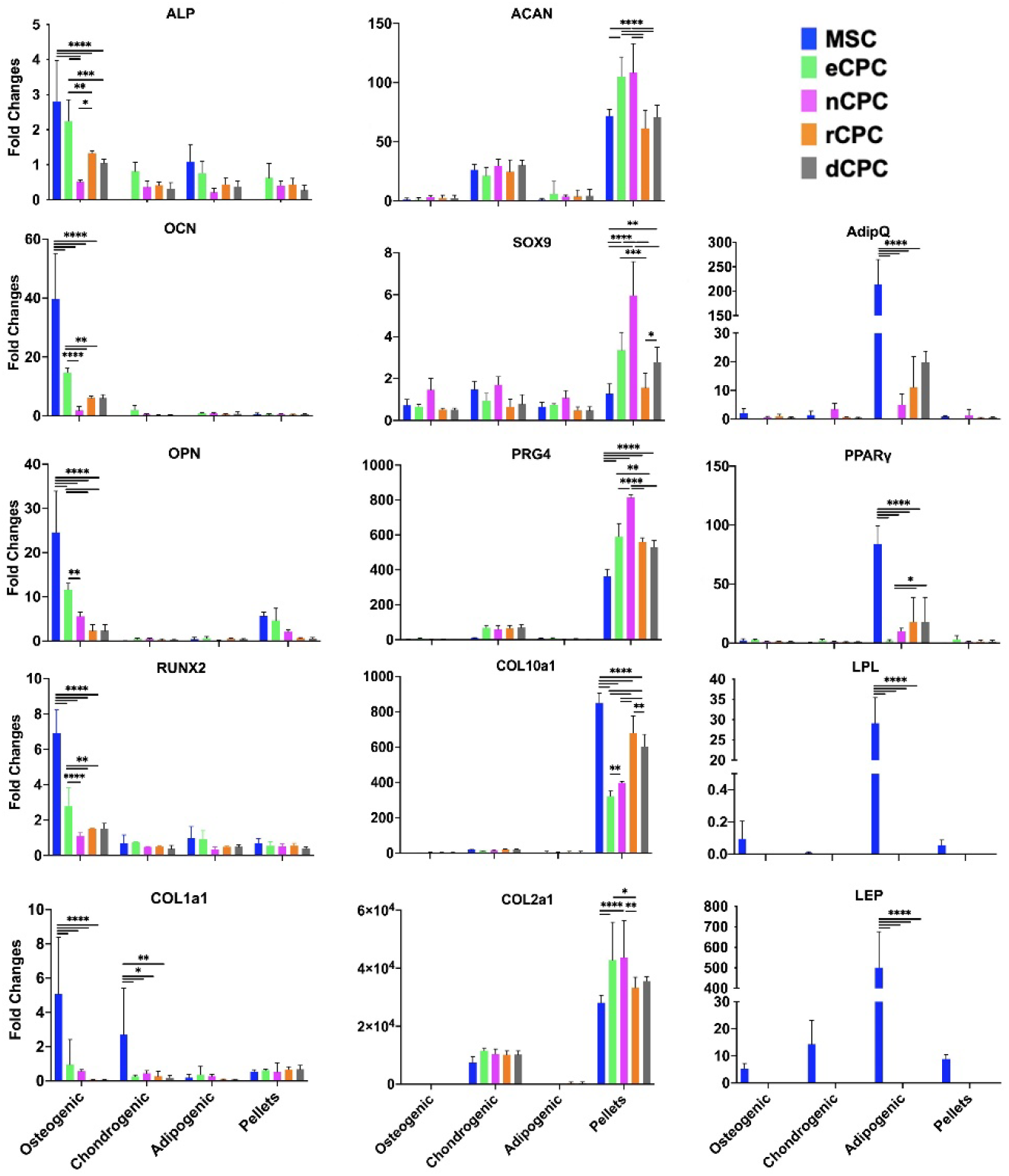
qRT-PCR of trilineage differentiated CPCs vs MSCs. The osteogenic (ALP, OCN, OPN, RUNX2, COL1a1), chondrogenic (ACAN, SOX9, PRG4, COL10A1, COL2A1), and adipogenic (AdipQ, PPAR γ, LPL, LEP) gene expression of MSCs and CPCs cultured in osteogenic, chondrogenic, or adipogenic medium in plate (2D) or in pellets (only for chondrogenesis) for 3 weeks. Results are reported as mean ± standard deviation of log2-fold change (delta delta CT method compared to the housekeeping gene and day 0), and statistical significance was denoted as *p < 0.05, **p < 0.01, ***p< 0.001, ****p < 0.0001 by a two way ANOVA followed by a Tukey post hoc multiple comparison analysis.

Chondrogenic gene expression was evaluated for both 2D plate culture and pellets (Figure 5). 2D culture yielded low chondrogenic gene expression among all cell types with no significant difference between any subset; however, when evaluated in the more appropriate pellet culture system, large differences were revealed. nCPCs and eCPCs showed a statistically significant upregulation of ACAN, SOX9, PRG4, and COL2a1 compared to MSCs, rCPCs, and dCPCs. This trend mirrors the observed differences in Alcian blue in Figure 4. Notably, the RNA levels of COL10a1 typical of a more hypertrophic phenotype were higher for MSCs, rCPCs, and dCPCs, compared to eCPCs, and nCPCs, with eCPCs having the statistically lower expression suggesting the least propensity towards hypertrophy. ^47^ Finally, CPCs exposed to the adipogenic medium expressed minimal adipogenic gene expression compared to the robust upregulation in MSCs, in accord with the negative Oil Red-O staining for CPCs in Figure 4.

### 3.4. CPCs display robust chondrogenesis and resist hypertrophy

CPCs and CCs response to hypertrophy was probed via a 14-day culture in chondrogenic medium to establish a chondral phenotype, followed by an additional 14 days in hypertrophic medium. After 28 days, deposition of sGAGs (histology), collagen II (COL II), collagen I (COL I), and collagen X (COL X) (immunohistochemistry) was assessed. Overall, 28 days of chondrogenesis resulted in a more robust sGAG and COL II deposition in eCPCs compared to corresponding eCCs. Exposure to hypertrophic medium, decreased slightly sGAGs and COL II deposition in both eCPCs and eCCs (Figure 6). However, eCCs exhibited stronger COL I and COL X staining in both chondrogenic and, more markedly, hypertrophic media than eCPCs.

**Figure 6.**
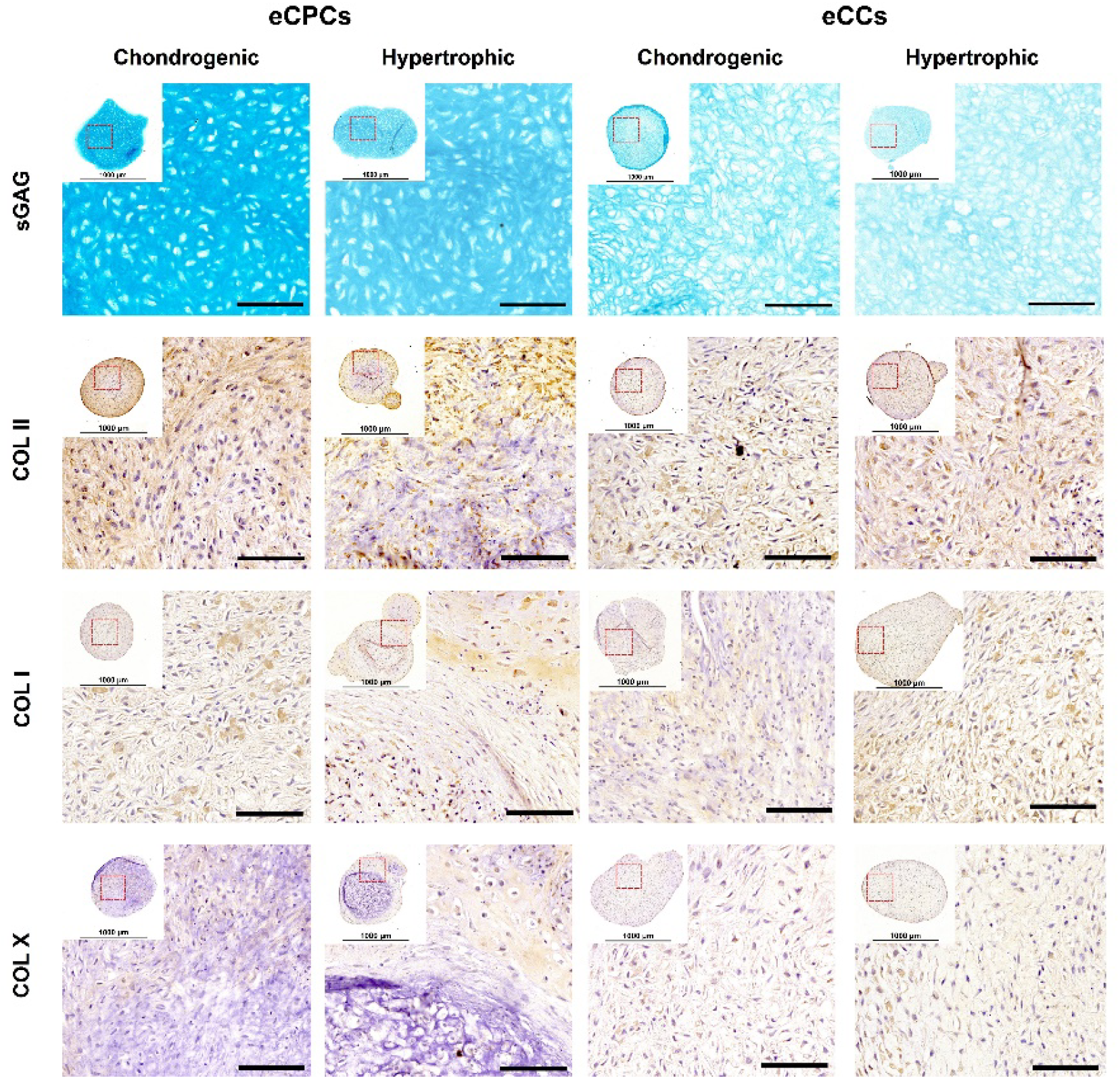
Immunohistochemistry of hypertrophic challenged eCPC pellets compared to eCCs. Alcian blue (row 1), collagen II (COL II row 2), collagen I (COL I row 3), and collagen X (COL X row 4) staining of eCPCs and eCCs pellets cultured in chondrogenic medium for 4 weeks (column 1 & 3) or chondrogenic then hypertrophic media for 4 weeks (2 weeks each, column 2 & 4). Scale bars = 100 μm.

Pellets of nCPCs and nCCs followed a similar trend to the eCPCs and eCCs in Figure 6. nCPCs showed higher sGAG and COL II deposition compared to nCCs in both chondrogenic and hypertrophic media, although with overall lower levels in the latter. COL I and X secretion was increased in nCCs compared to nCPCs under both chondrogenic and hypertrophic conditions (Figure 7). Finally, when this 28 day study was conducted with both rib and joint cells, a decrease in COL II and sGAGs along with an increase in COL I or COL X was observed under hypertrophy compared to the chondrogenic control, with CPCs generally having a lesser hypertrophic matrix deposition response than chondrocytes (Figure 8 & 9).

**Figure 7.**
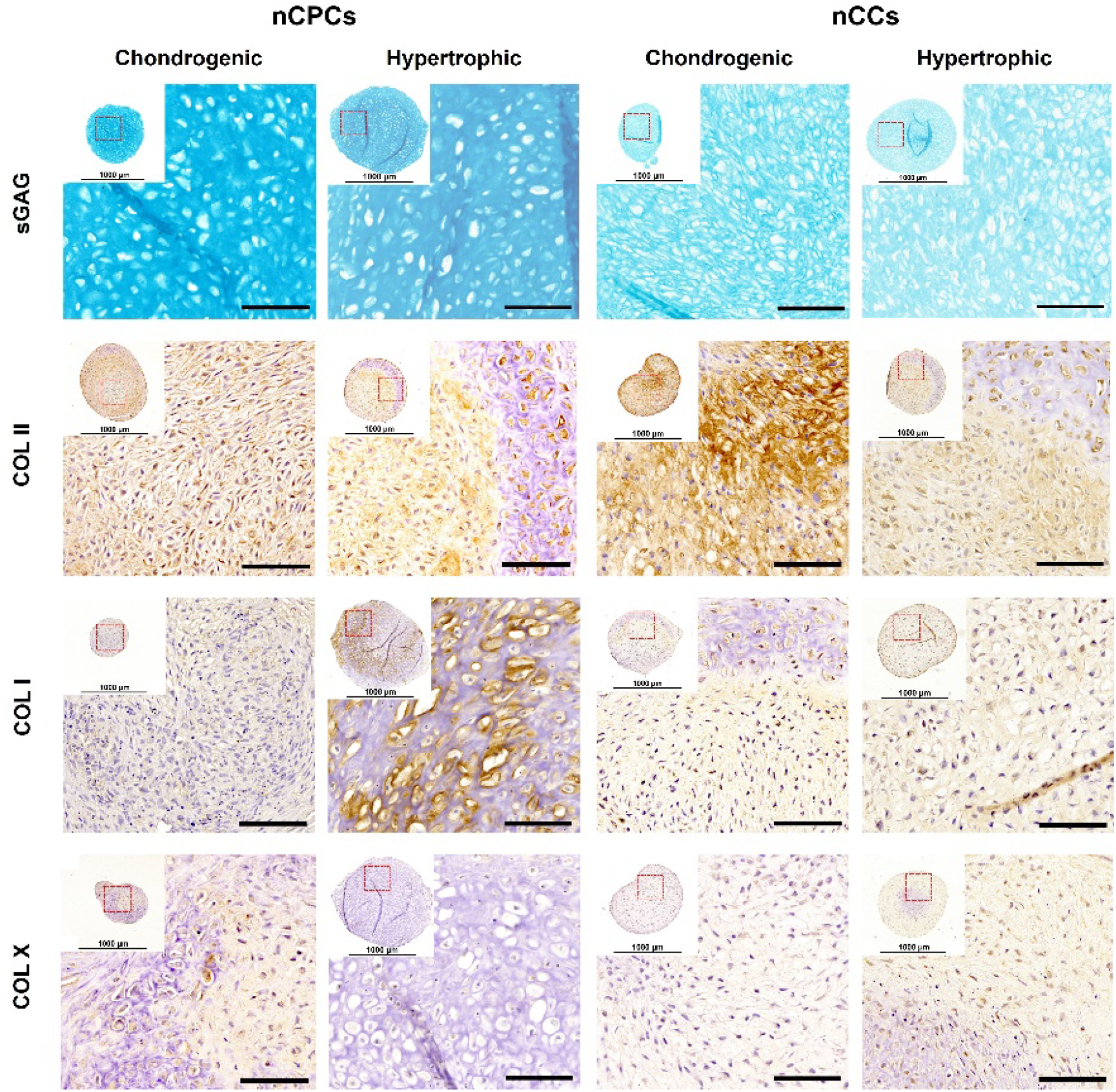
Immunohistochemistry of hypertrophic challenged nCPC pellets compared to nCCs. Alcian blue (row 1), collagen II (COL II row 2), collagen I (COL I row 3), and collagen X (COL X row 4) staining of nCPCs and nCCs cultured in chondrogenic medium for 4 weeks (column 1 & 3) or chondrogenic then hypertrophic media for 4 weeks (2 weeks each, column 2 & 4). Scale bars = 100 μm.

**Figure 8.**
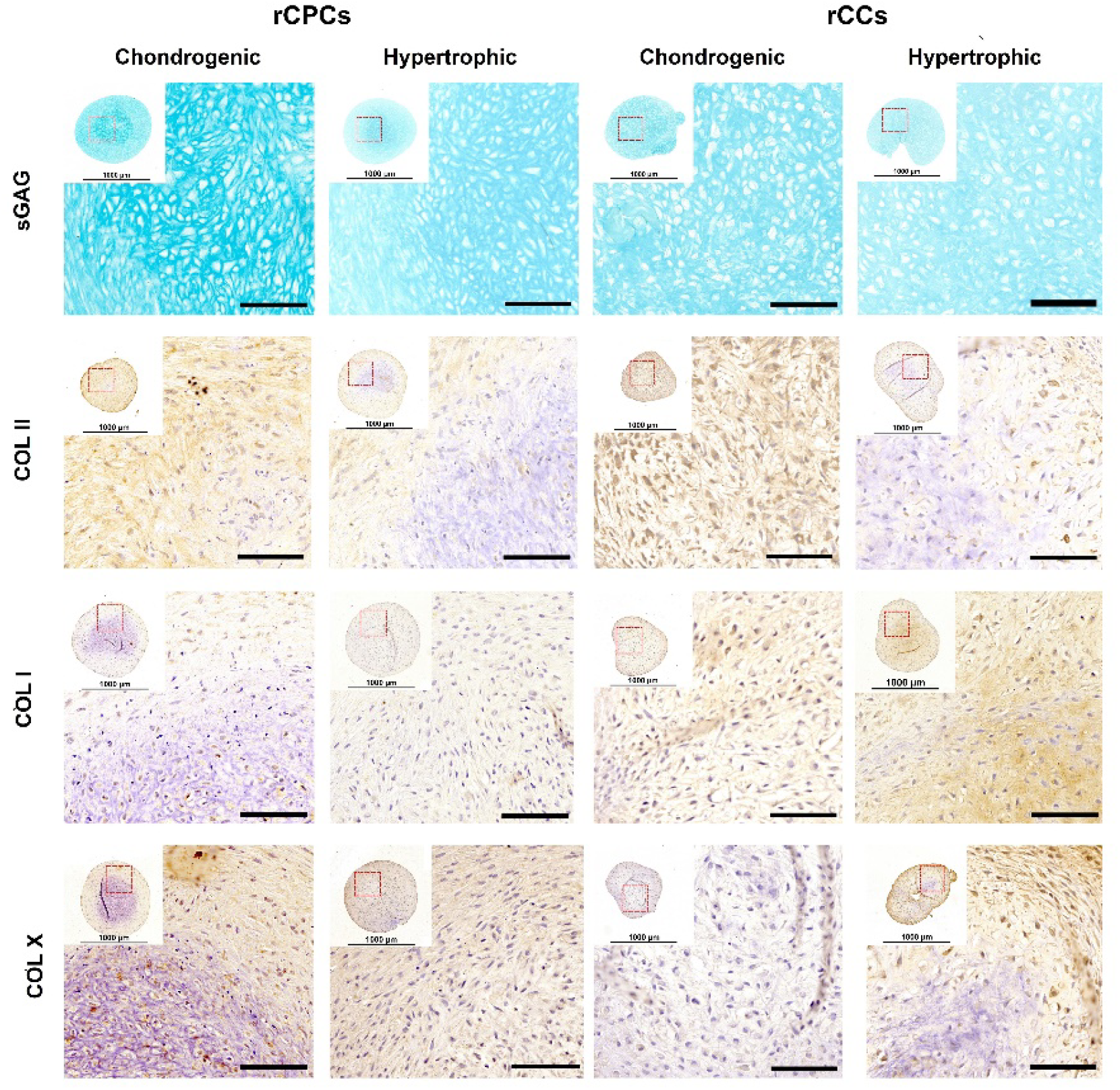
Immunohistochemistry of hypertrophic challenged rCPC pellets compared to rCCs. Alcian blue (row 1), collagen II (COL II row 2), collagen I (COL I row 3), and collagen X (COL X row 4) staining of rCPCs and rCCs cultured in chondrogenic medium for 4 weeks (column 1 & 3) or chondrogenic then hypertrophic media for 4 weeks (2 weeks each, column 2 & 4). Scale bars = 100 μm.

**Figure 9.**
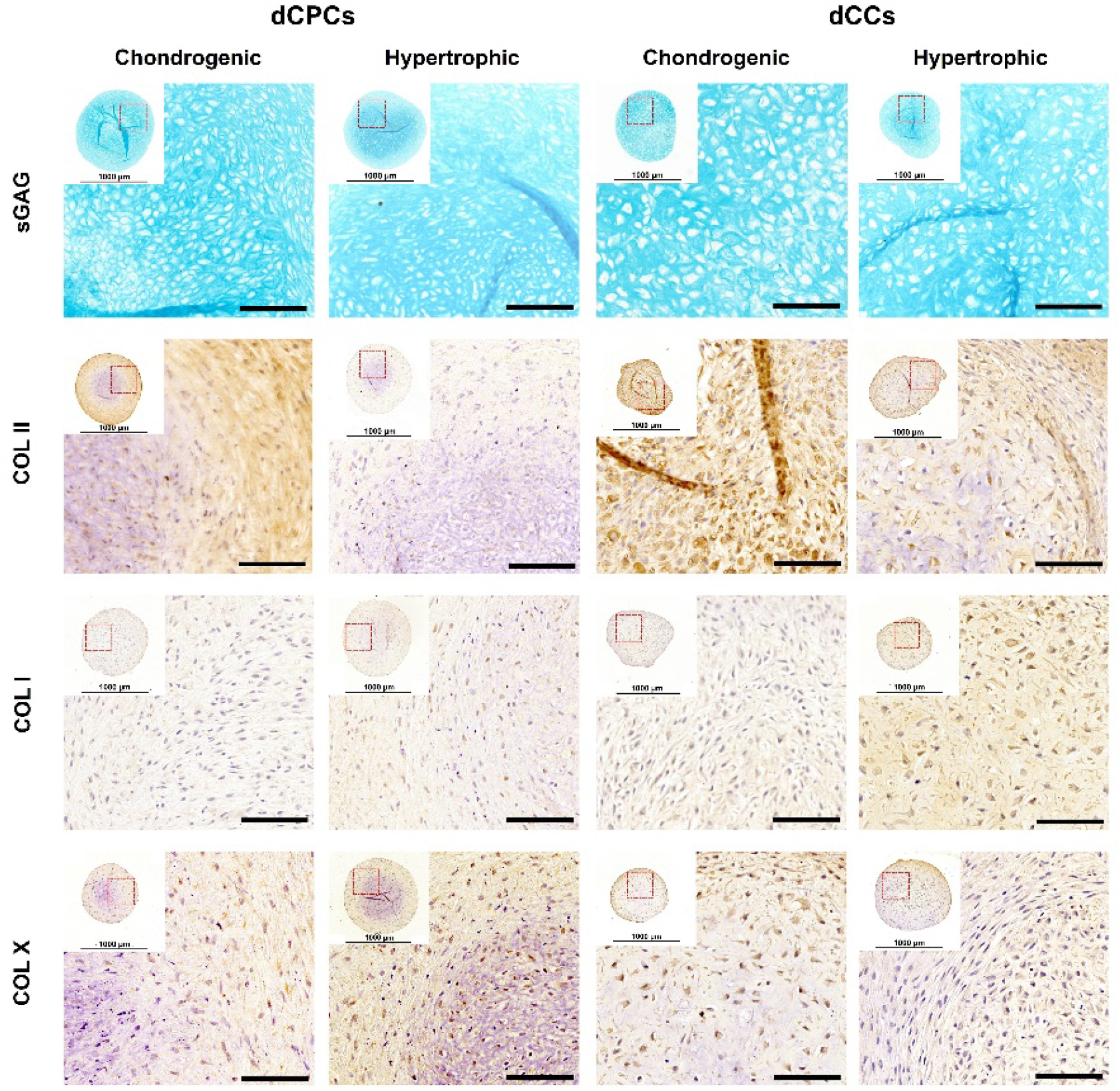
Immunohistochemistry of hypertrophic challenged rCPC pellets compared to rCCs. Alcian blue (row 1), collagen II (COL II row 2), collagen I (COL I row 3), and collagen X (COL X row 4) staining of dCPCs and dCCs cultured in chondrogenic medium for 4 weeks (column 1 & 3) or chondrogenic then hypertrophic media for 4 weeks (2 weeks each, column 2 & 4). Scale bars = 100 μm.

To quantitatively analyze the matrix compositions of the chondrogenic and hypertrophic pellets, the GAG and collagen content for each cell type was measured and is reported as a ratio between day 28 and day 14 (Figure 10). The full content and weight of GAGs at day 14, 21, and 28 is reported in the supplemental section (Figure S2).

**Figure 10.**
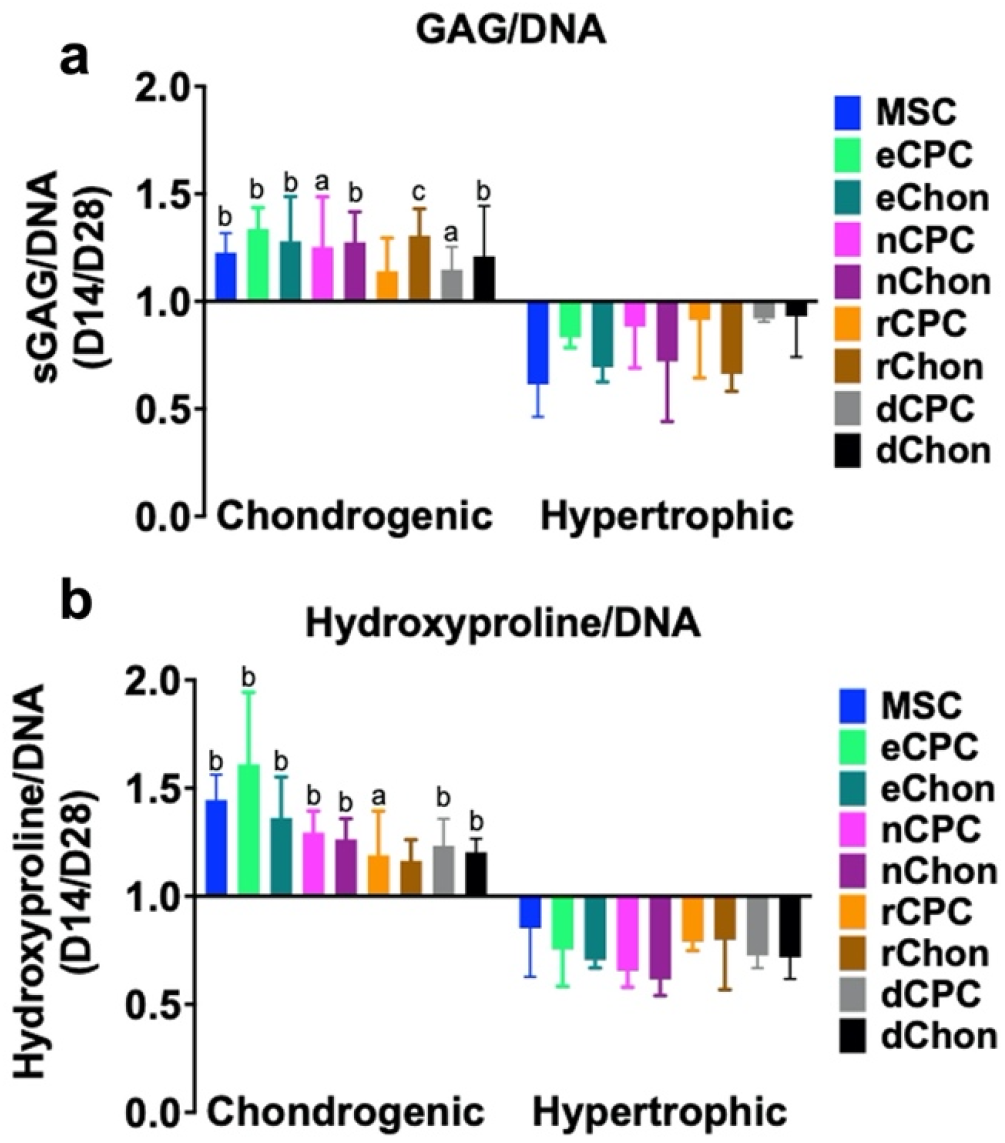
GAG and DNA quantification following 2 weeks of hypertrophic induction. (A) The GAG/DNA and (B) Collagen/DNA quantification of MSCs, CPCs or CCs pellets after chondrogenic or hypertrophic treatment for 28 days normalized to GAG/DNA or Collagen/DNA content at day 14. Results reported as mean ± standard deviation with statistical significance denoted by letter a = p<0.05, b = p<0.01, c = p<0.001 as compared to the corresponding hypertrophic condition using a paired students’ t-test.

Under the chondrogenic medium condition, all pellets showed an increase in GAG and hydroxyproline content on day 28 compared to day 14, indicative of a retained chondrogenic phenotype. In contrast, both the GAG (Figure 10a) and collagen content (Figure 10b) decreased significantly after 14 days of hypertrophic treatment suggesting that the lower signals for sGAGs and COL II in Figures 6-9 is the result of matrix remodeling and degradation. GAG loss under hypertrophic condition was most significant in MSCs and CCs, whereas the hydroxyproline changes did not follow any specific trend. Notably, the assay is insensitive to specific collagen types, thus suggesting that the observed decreases in COL II in Figures 6-9 were likely offset by the hypertrophic increase in COL I and COL X.

Lastly, pellets were evaluated using a panel of chondrogenic (*COL2A1, PRG4, COMP*, and *ACAN*) and hypertrophic/osteogenic (*RUNX2, BMP2, BMPR1A, BMPR1B, BMPR2, MMP13, IHH, COL10A1, ALP*, and *ACVR1*) associated genes. Genes involved in osteogenesis were expressed significantly higher by MSCs compared to any CPC. Some hypertrophic/osteogenic associated genes were also more highly expressed by CCs in the hypertrophic media compared to the corresponding CPC subset. This lesser increase in expression by CPCs point to a stable chondral phenotype and suggests some resistance to hypertrophy and ossification. Furthermore, nose and ear CPCs’ higher *BMPR1B* and *BMPR2* expression, especially in hypertrophic conditions, may suggest initial lower levels of those receptors for BMP signaling, a hypertrophic driver, consistently with eCPCs and nCPCs lesser response in Figure 6-9. *IHH*, an early hypertrophy marker, showed an increased expression in dCCs, dCPCs, rCCs, and rCPCs while it was negligible in eCPCs, eCCs, nCPCs, and nCCs. *ALP* and *ACVR1* are early osteogenic markers that were increased in hypertrophic medium and were markedly higher in MSCs and chondrocytes rather than CPCs. This same tendency was observed for the osteogenic matrix gene *COL1A1* and for the hypertrophic marker *COL10A1* for rib and digit cells, but not for ear and nose, although *COL10A1* increased for nCCs under hypertrophy. *MMP13* serves to degrade and remodel cartilage and its gene expression increased in hypertrophy, especially for MSCs, rCCs, and dCPCs, whereas eCPCs and nCPCs were not affected by the hypertrophic medium. Finally, chondrogenic genes such as *COL2A1, PRG4, COMP*, and *ACAN* were generally expressed at higher levels in chondrogenic media rather than the hypertrophic condition. A notable exception is the expression of *ACAN* for ear and nose cells that remains high even under hypertrophy.

## 4. Discussion

CPCs represent a major milestone in understanding cartilage repair. Harnessing these subset of progenitor cells especially from sites that are easily accessible and minimally invasive marks a significant opportunity for therapeutic relevance. To date, there existed no comparisons between CPCs isolated from human ear, nose, rib, and joints. Our unique location inside the Children’s Hospital of Philadelphia endowed us a streamlined process for cartilage collection and CPC extraction and allowed for a statistically significant number of donors to be collected for each cartilage type. In this study, we focused on the robust and extensive characterization of these CPC subtypes compared to both MSCs and the corresponding tissue CCs through the comparative analysis of proliferation, differentiation, and hypertrophic stress response. We expect that this work will help identify the most promising progenitor cells sources in terms of accessibility and performance for cartilage repair.

Prior to the emergence of CPCs, MSCs were broadly considered the gold standard for cartilage repair, notwithstanding their well-known limitations: the propensity to undergo hypertrophy and calcify especially *in vivo*, a less robust and stable chondral phenotype than chondrocytes, and the generation of a more fibrous cartilage with relatively high collagen I. These limitations were also confirmed in our study. Tri-lineage differentiation evidenced how MSCs can acquire an osteogenic, adipogenic, or chondrogenic phenotypes, whereas all CPC subtypes could only undergo chondrogenesis and exhibited limited markers of osteogenesis and adipogenesis (Fig. 4-5). While the ability of MSCs to differentiate into 3 lineages allows them to be used for the treatment of multiple tissues, we observe that their lineage specification is not always robust, with relatively high expressions of the osteogenic/fibrous gene *COL1* and the hypertrophic marker *COL10* even during chondrogenic differentiation.^48^ The development of a more fibrocartilaginous phenotype (*COL1*) is well-documented in numerous studies for MSC differentiation,^48,49^ and while potentially useful for meniscus tissue engineering,^50^ it is sub-optimal for hyaline cartilage repair. This is a well-known challenge in the use of MSCs as is their unstable chondrogenic phenotype prone to calcification (*COL10*) even without the addition of hypertrophic inducing agents. Notably, *COL1* is not upregulated in any CPC subtype during chondrogenesis and *COL10* is less upregulated than in MSCs, with ear CPCs exhibiting the least *COL10* increase of all compared CPC types (Fig.5). This suggests the induction of a more stable hyaline phenotype than that of MSCs. To further confirm these limitations of MSCs, we induced hypertrophy via the incorporation of triiodothyronine (T3) into the differentiation medium of pellets after two weeks of chondrogenic pre-differentiation. T3 is a powerful morphogen that has been shown to induce hypertrophic terminal differentiation via the initial upregulation of BMP-4 signaling ultimately leading to matrix calcification.^12,51,52^ When exposed to T3, MSCs have been reported to exhibit a marked shift in gene expression for *COL2* to *COL1* and a substantial upregulation of *COLX*, the hallmark of chondrocyte hypertrophy.^52^ By harnessing this well-defined induction pathway, we challenged our MSC-derived chondral pellets (Fig.11, blue bar columns). Congruent with the literature, MSCs displayed substantial hypertrophic potential through increased upregulation of *ALP, COLX, BMP2*, and *BMPR2*, and through a shift from *COL2A1* (downregulated) to *COL1A1* (upregulated) compared to the non-T3 chondrogenic control (Fig. 11). These observations are congruent with MSCs hypertrophy studies.^12^ Notably, this shift in gene expression was significantly more pronounced than any CPC subtype, all of which appeared to resist the hypertrophic challenge to a greater extent than MSCs. In agreement with these results, GAG and collagen matrix deposition in MSC pellets were decreased substantially, albeit not differently from CPCs, following 14 days of hypertrophy suggesting remodeling of the cartilaginous matrix (Fig. 10).

**Figure 11.**
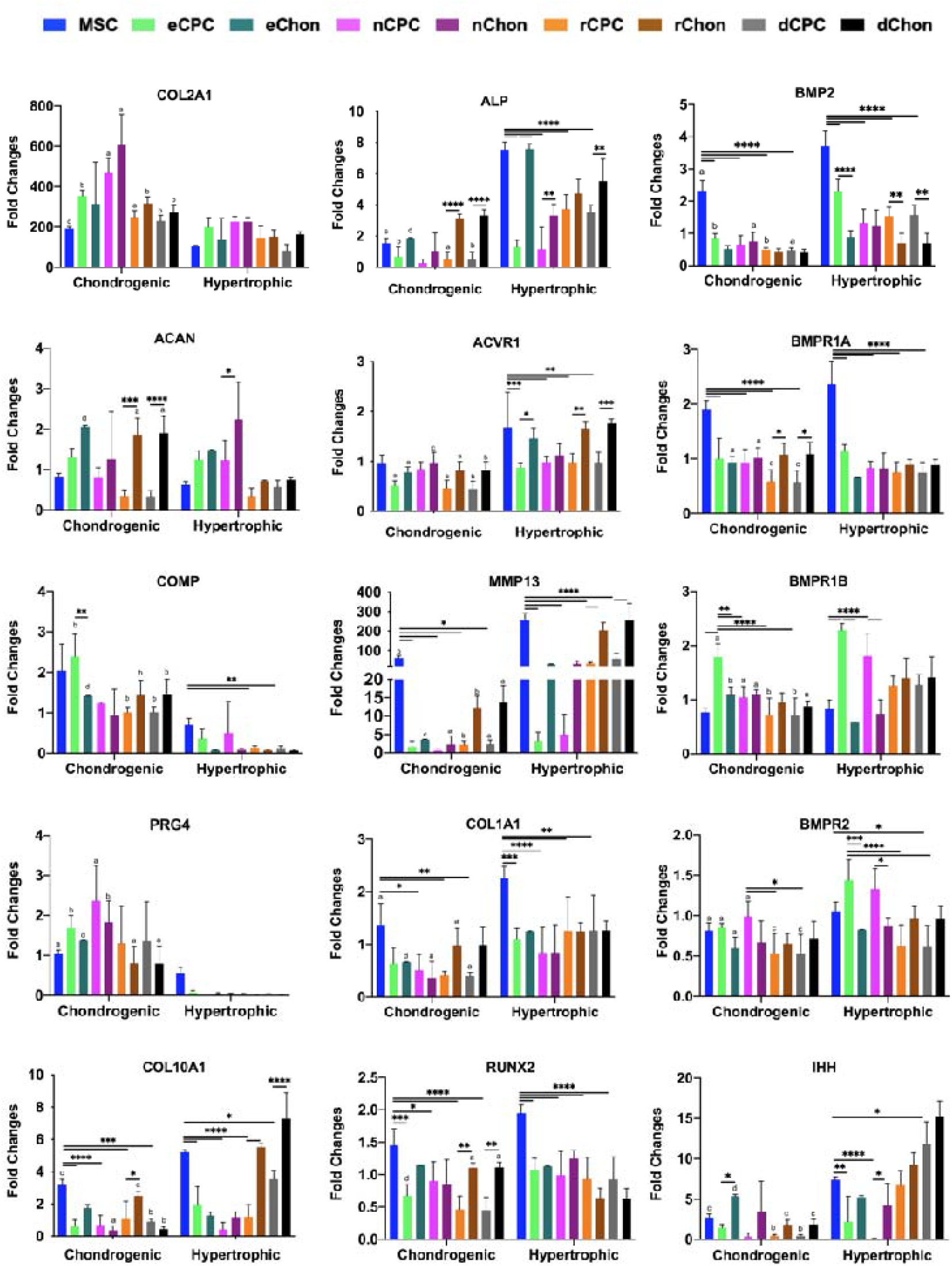
Gene expression of chondrogenic or hypertrophic induced pellets. Chondrogenic (COL2A1, PRG4, COMP, and ACAN) and hypertrophic/osteogenic (RUNX2, BMP2, BMPR1A, BMPR1B, BMPR2, MMP13, IHH, COL10A1, ALP, and ACVR1) associated genes of MSCs, CPCs and CCs pellets cultured with chondrogenic medium for 4 weeks (to the left on each graph) or chondrogenic medium for 2 weeks followed by hypertrophic media for 2 weeks (4 weeks total, to the right on each graph). Results reported as mean ± standard deviation of log2-fold change of week 4 vs. week 2, and statistical significance was denotated as (a,*) for p < 0.05, (b,**) p < 0.01, (c,***) p< 0.001, (d,****) p < 0.0001 by a two way ANOVA followed by a Tukey post hoc multiple comparison analysis, with letters comparing the same cell type between chondrogenic vs hypertrophic condition, and asterisks comparing between cell types within the same condition.

Besides MSCs, cartilage chondrocytes (CCs) have also been widely used for tissue engineering and cartilage repair, with matrix-induced autologous chondrocyte implantation (MACI) as one such example. MACI has recently been FDA approved but prior to that it had already been used for years in Europe and correlated with some of the most positive outcomes in hyaline cartilage repair. However, it has been reported that nearly 25% of MACI implants do not fully restore joint function and some become fibrocartilage.^53,54^ Key translational issues with CCs are their slow proliferation, de-differentiation during expansion, and donor site morbidity at the site of harvest in the joint. To overcome some these pitfalls, the use of autologous nasal chondrocytes cultured with phenotypic stabilizers (FGF-2, TGF-β1, and BMP inhibitors) has been proposed. This “nose-to-knee” approach has shown initial results in human that are potentially superior to MACI^55^ so much as to warrant a full clinical trial. Notably, their study, Mumme *et al*.,^55^ was able to limit the de-differentiation of CCs, but still required a 4-week *in vitro* expansion. In our present work, we directly compared proliferation of CCs and CPCs extracted from the same cartilage subtype samples of human ear, nose, rib, and joint of multiple donors, as well as of MSCs. As reported extensively in the literature, all CC subtypes displayed a substantially slower proliferation rate compared to MSCs; however, CPCs not only proliferated faster than the corresponding CCs, but also of MSCs, making CPCs relevant for translational approaches (Fig. 3). Nonetheless, proliferation rate is not the only challenge in the use of CCs for cartilage repair.

Chondrocytes may undergo terminal differentiation via hypertrophy and calcification similarly to what happens during development^3,48^ and this is one of the most common failure modes of cartilage repair or tissue engineering using chondrocytes.^56^ We compared all CCs and paired CPC subtypes in terms of chondrogenic maturation (4 weeks) and response to a hypertrophic challenge by T3 (2 weeks of maturation + 2 weeks of T3 challenge). After 4 weeks of chondrogenesis, all CPCs present a robust Alcian blue staining that was comparable or better (eCPCs and nCPCs) than the corresponding CCs (Fig. 6-9, first and third columns), suggesting a similar or even better GAG deposition for CPCs, a trend supported by the GAG/DNA quantification (Fig. S2). In terms of collagen II, collagen I and collagen X, similar levels of staining are observed for both CCs and CPCs, with only nCCs having higher levels of collagen II than the corresponding nCPCs, and rCCs having lower levels of collagen X than the corresponding rCPCs (Fig. 6-9, first and third columns). This confirms the high performance of all CPC subtypes in terms of their chondrogenic potential which is equivalent to that of CCs. Following hypertrophic challenge, there was an overall decrease in collagen II content and increase in collagen I and collagen X, accompanied by a more aligned and fibrous matrix organization. These changes in composition and structure are likely associated with terminal differentiation and the initiation of the remodeling associated with calcification. Although each CC subtype displayed varying initial levels of matrix deposition, the trends of changes associated with T3 challenge remained consistent across each subtype (Fig. 6-9, third and fourth columns). This is especially interesting considering that chondrocytes from ear and nose do not naturally undergo endochondral ossification during development; yet, ear and nose chondrocyte populations still responded to T3 suggesting they do not have an intrinsic resistance to hypertrophy and calcification. Gene expression analysis confirmed these trends for CCs, with a general decrease in pro-chondrogenic genes and an increase in calcification and remodeling genes. Interestingly, rCCs and dCCs sourced from cartilage that undergoes endochondral ossification, presented the highest expression of *COL10A1* (hypertrophy), *IHH* (osteogenesis/calcification), and *MMP13* (remodeling) when challenged with T3. While this might suggest tissue-specific memory for chondrocytes, a definitive conclusion cannot be made as our samples are derived from pre-adolescent donors, and these findings should be confirmed in tissues from adult, skeletally mature sources. As for CPCs, when challenged with T3, each subtype followed the same histological decrease in collagen 2 and increase in collagen 1; notably, however, collagen X deposition was not present following hypertrophic conditioning. The absence of secreted collagen X, the hallmark of hypertrophy and terminal differentiation during endochondral ossification, may suggest that CPCs of each subtype do not undergo terminal differentiation as readily as CCs. Gene expression for each CPCs subtype confirmed decreases in *COL2, COMP, PRG4* and increases in *COL1, ALP, ACVR1, BPR1B*, and *BMPR2* expression. However, CPCs did not show a significant increase in *COLX* after hypertrophic challenge except for dCPCs, while only eCPCs and nCPCs did not show an increase in *IHH* and *MMP13*. In general, CPCs responded less to the T3 challenge than the corresponding CCs, suggesting they might be further away from terminal differentiation than primary chondrocytes. Finally, MSCs displayed overall the most pronounced expression for hypertrophic/calcification genes both at baseline after chondrogenesis and following challenge with T3. Taken together these observations suggest that CPC, especially from ear and nose, might have the lowest risk for calcification.

Our results strongly point to CPCs as a cell source potentially superior to both MSCs and CCs for cartilage repair. Strikingly, CPCs proliferation rate is nearly 2-fold greater than MSCs and over 4-fold greater than CCs. This could be attributed to higher colony formation ability in CPCs as a CD56^+^ population (Fig. 2) compared to CD56^-^ MSCs ^26^. Therapeutic strategies could significantly benefit from CPCs’ high proliferative capacity which allows for a shorter translational timeline. For instance, any time that chondrocytes are employed in cartilage repair, a simple fibronectin selection could isolate the CPCs resulting in faster availability to obtain a high number of cells for translation. The limited osteogenic and negligible adipogenic capacity suggests that CPCs are further ahead in terms of lineage commitment than MSCs reinforcing their identity as committed chondrocyte precursor subset^57^. Among the different CPC sources, trilineage differentiation points to the ear and nose CPC as the most promising, with the strongest Alcian blue staining (Fig. 3), the highest upregulation of *COL2a1, ACAN, PRG4* and SOX9, and a lower baseline expression of *COL10a1* compared to the other subsets and MSCs (Fig. 5). Under hypertrophic challenge, all CPCs showed equal or superior resistance compared to the corresponding chondrocyte population. For all CPCs subtypes there was small (nCPC and rCPC) to no increase (eCPC and rCPC) in collagen I or collagen X deposition compared to the increases displayed by corresponding CCs. Moreover, no CPC subset visually lost collagen II suggesting that while CPCs may secrete collagen I during the first 2 weeks of hypertrophic challenge, they do not remodel and breakdown the existing cartilaginous environment. This is supported by the gene expression changes after T3 challenge, where *MMP13* expression associated with collagen remodeling is significantly upregulated in CCs compared to CPCs as is the expression of *ALP* and *ACVR1* both involved in the substitution of cartilage with bone during endochondral ossification. Interestingly, the two most chondrogenic CPC types, eCPCs and nCPCs, are found in cartilage types that do not normally undergo endochondral ossification. Conveniently, these superior chondrogenic CPCs are in regions of the body, ear and nose, that are most easily accessible for minimally invasive biopsies that can be achieved in a cosmetically invisible manner.^24,55^ However, only recently eCPCs have been explored for cartilage tissue engineering by Otto *et al*. who found superior mechanical properties and GAG deposition for constructs made of eCPCs encapsulated in a methacrylated gelatin hydrogel.^17^ While nCPCs performed similar to eCPCs, their existence and properties aside from our characterizations have only been sparsely reported^7^.

In conclusion CPCs are a highly proliferative and chondrogenic cell type that resists hypertrophy and calcification. Compared to both MSCs and CCs, CPCs may then represent a more translatable and reliable cell source for cartilage repair. While each CPC subtype’s properties differ by cartilage source, all CPCs displayed similar surface markers, differentiation capacities, and gene expression. Of the CPC subsets explored, ear and nose CPCs were generally superior to joint and rib CPCs. Both ear and nose CPCs had the highest chondrogenic gene expression and matrix secretion while resisting hypertrophy the most in both histology and gene expression. For tissue engineering, CPCs can be rapidly expanded reaching within 10 days a sufficient number to be seeded into a scaffold for cartilage repair while avoiding the challenges associated with de-differentiation of chondrocytes during *in vitro* expansion or the more invasive nature of MSCs harvest. The easily accessible nature of ear and nose CPCs presents significant opportunities for translational approaches with a shortened turnaround time from cell harvest to implantation. Overall, CPCs combine all of the most desirable properties of rapid proliferation, stable differentiation, and ease of access that make them an ideal choice for translational cartilage repair.

## 5. Contributors

RG, SA, and BJ conceived the study and planned the experiments. RG, SA, PG, and TG performed the experiments. RG, SA, PG, and KS analyzed the data. RG, SA, and PG prepared the manuscript. All authors approved the finalized version of the manuscript.

## 7. Acknowledgments

We thank the laboratory of Dr. Véronique Lefebvre and specifically Dr. Marco Angelozzi for their histological facilities and slide scanner. We also thank Dr. John Germiller, Dr. Ian Jacobs, and Dr. Jesse Taylor for the collected cartilage tissues. We acknowledge Dr. Brian Johnstone and Mr. Ajit Elhance for providing protocols for cell maintenance and biochemical assays. We thank the laboratory of Dr. Evangelia Bellas for the kind gift of adipogenic medium. We acknowledge Children’s Hospital of Philadelphia’s Flow Cytometry Core Laboratory and Pathology Core Laboratory for their respective services. We thank the Children’s Hospital of Philadelphia’s Center for Applied Genomics and specifically Dr. Renata Pellegrino for aiding with the design of Fluidgm PCR array.

This work was supported in part by the National Heart, Lung, and Blood Institute of the National Institute of Health (R21HL159521 to R.G.), the Children’s Hospital of Philadelphia Research Institute (R.G.), the Frontier Program in Airway Disorders of the Children’s Hospital of Philadelphia (R.G.), and the National Science Foundation Graduate Research Fellowship No. DGE 1845298 (M.R.A. and P.M.G.).

## 10. Supplementary

**Figure S1.**
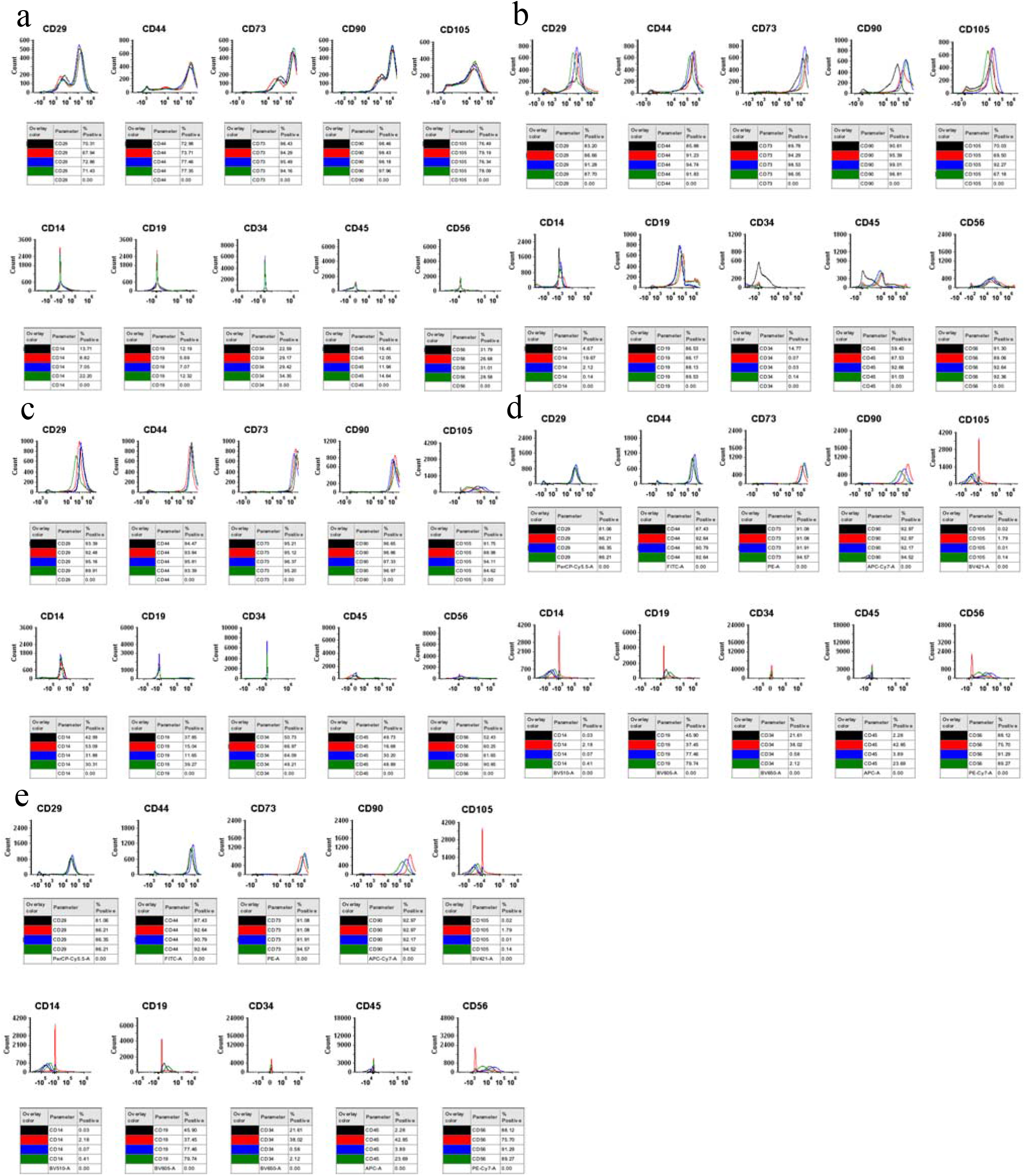
Immunophenotyping of CPCs and MSCs. Flow cytometry immunophenotyping of (a) MSCs, (b) eCPCs, (c) nCPCs, (d) rCPCs, (e) dCPCs. Histograms of CD29, CD44, CD73, CD90, CD105, CD14, CD19, CD34, CD45, CD56 at passage 3 were reported.

**Figure S2.**
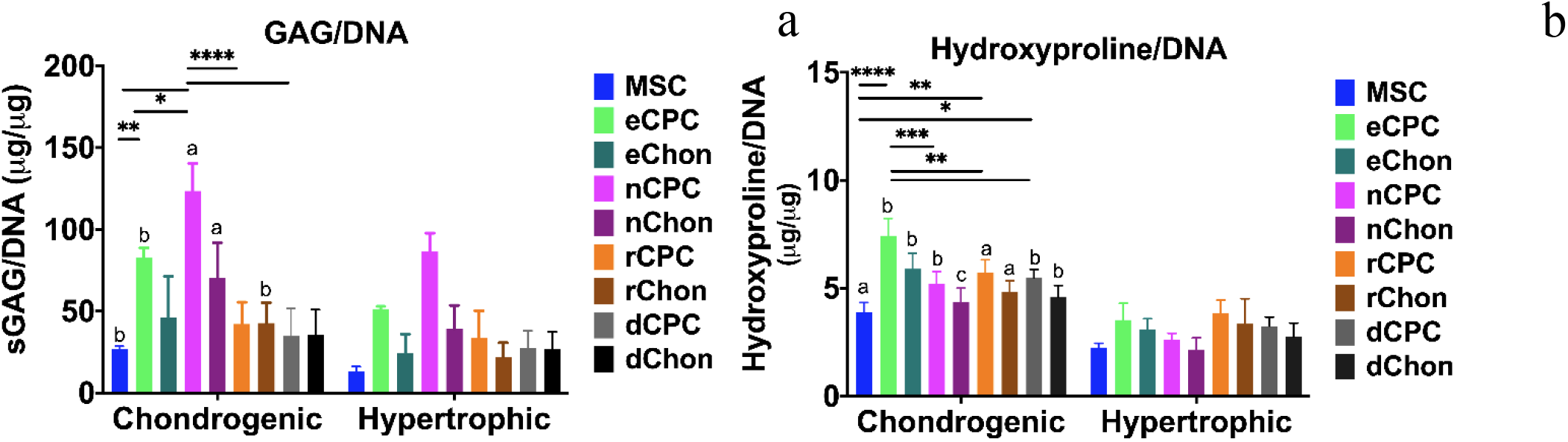
Total GAGs and hydroxyproline content normalized to DNA for the chondrogenic and hypertrophic induced pellets. (a) GAGs or (b) hydroxyproline dry weight per μg of DNA for MSCs, CPCs or CCs pellets after chondrogenic or hypertrophic treatment for 28 days. Results reported as mean ± standard deviation. Statistical significance denotated as (a,*) for p < 0.05, (b,**) p < 0.01, (c,***) p< 0.001, (d,****) p < 0.0001 by a two way ANOVA followed by a Tukey post hoc multiple comparison analysis, with letters comparing the same cell type between chondrogenic vs hypertrophic condition, and asterisks comparing between cell types within the same condition.

